# A Domain-Specific Language Model Approach for Identifying Immunogenic Epitopes from Publicly Available Data: EpitopeMiner

**DOI:** 10.1101/2025.01.21.633814

**Authors:** Wei Kit Tan, Wei Lin Tang, Agamjyot Singh Chadha, Roopa Niedu, Wendy Lee, Jing Quan Lim, Menaka Priyadharsani Rajapakse, Solomon Wilson, Kaibo Duan, Raymond Wei Liang Chong, Choon Kiat Ong, Bernett Lee, Olaf Rotzschke, Mai Chan Lau

**Affiliations:** Bioinformatics Institute (BII), Agency of Science, Technology and Research (A*STAR), 30 Biopolis Street, Matrix Building, Singapore 138671, Republic of Singapore; Singapore Immunology Network (SIgN), Agency of Science, Technology and Research (A*STAR), 8A Biomedical Grove, Immunos Building, Singapore 138668, Republic of Singapore; Lymphoma Translational Research Laboratory, Division of Cellular and Molecular Research, National Cancer Centre Singapore, 30 Hospital Boulevard, Singapore 168583, Republic of Singapore; ONCO-ACP, Duke-NUS Medical School, 8 College Road, Singapore 169857, Republic of Singapore; Lee Kong Chian School of Medicine (LKCMedicine), Nanyang Technological University Singapore, Singapore 636921, Republic of Singapore; Cancer and Stem Cell Biology, Duke-NUS Graduate Medical School, 8 College Road, Singapore 169857, Republic of Singapore

## Abstract

Personalized cancer vaccines represent a transformative approach in immunotherapy, leveraging tumor-specific antigens, such as neoantigens, to stimulate durable and targeted immune responses. However, identifying immunogenic neoantigen epitopes through conventional approaches - relying on in silico predictions followed by experimental validation - remains a significant challenge due to limitations of computational tools and scattering of experiment data. To address these challenges, we introduce EpitopeMiner, a domain-specific large language model (LLM) enhanced with a Retrieval-Augmented Generation (RAG) framework, specifically designed to identify immunogenic epitopes predicted by existing computational tools. EpitopeMiner utilizes a custom database of MHC-I-associated epitope-related literature to provide domain-specific knowledge, enhancing the precision and relevance of LLM-generated responses. EpitopeMiner offers three key features: First, the ability to identify epitopes with similar sequences and potentially similar immunogenic effects, which is especially valuable for neoantigens that are patient-specific and rarely found in public datasets. Second, it supports multiple epitope searches with structured outputs, enhancing scalability. Third, it provides original text chunks and paper identifiers, significantly simplifying validation and further exploration of the retrieved knowledge. Applying EpitopeMiner to well-characterized MHC class I epitopes demonstrates its ability to consistently retrieve relevant papers, efficiently providing targeted insights on T-cell response and immunogenicity, outperforming a commercial AI-powered literature tool. Notably, when applied to lymphoma patient-derived neoantigens, EpitopeMiner successfully retrieves immune response information for a few similar epitopes, despite operating with a relatively smaller database compared to the benchmark tool. All in all, EpitopeMiner bridges the gap between computational prediction and experimental validation, providing a scalable solution for extracting knowledge from public data, fostering cross-study synergies, and accelerating the development of personalized cancer vaccines.

## Introduction

Personalized cancer vaccines, a crucial component of immunotherapy in neoadjuvant or adjuvant settings, show great potential in inducing specific and robust immune responses, leading to effective tumor clearance(1-3). These cancer vaccines introduce tumor-specific antigens, such as neoantigens derived from a patient’s tumor mutations, to stimulate the immune system. These neoantigens are processed into peptides that bind to major histocompatibility complex (MHC) molecules, forming peptide-MHC complexes. These complexes are then presented on the surface of antigen-presenting cells (APCs) to CD8+ cytotoxic T cells. Through T-cell receptor (TCR) recognition, the immune system identifies and targets cancer cells, activating a robust and specific immune response to eliminate tumors(4).

While there is increasing recognition of the importance of antigen processing (e.g., NetChop(5)), peptide transport (e.g., NetCTL(6)), and immunogenicity (e.g., Immune Epitope Database (IEDB) analysis resource (AR) tools such as Immunogenicity and Deimmunization(7, 8)), most efforts remain focused on modeling peptide binding to MHC-I molecules, with widely used tools such as NetMHCpan(9) and MHCflurry(10). Additionally, there is no established consensus for setting cutoffs within individual in silico methods or across different tools, further complicating their application.

Advancements in experimental techniques, particularly mass spectrometry (MS) immunopeptidomics, have shown that although in silico predictions may identify thousands of somatic mutations and a few hundred potential MHC binders, the majority of these neoepitopes are not detected in tumors. Furthermore, using MHC–peptide tetramer staining to assess tumor-infiltrating lymphocytes (TILs), only a small subset of neoantigens is found to elicit T-cell responses. However, these measurement techniques also have limitations(11). MHC–peptide tetramer staining often requires ex vivo TIL expansion in ‘cold’ tumors, potentially altering T-cell specificities. Similarly, MS-based immunopeptidomics faces limited sensitivity for identifying naturally presented MHC ligands, relies on costly equipment, and demands substantial cell quantities, making it technically challenging.

In-silico predicted epitope candidates undoubtedly require experimental validation (12). Given the limitations and costs involved, establishing an effective knowledge-sharing framework is crucial. Tens of thousands of peptides—eluted from MHCs, identified by MS, and some tested via MHC–peptide tetramer staining—exist scattered across literature, public databases such as IEDB(8), dbPepNeo(13) and NeoDB(14), supplementary data in papers, and even laboratory records (11). Reviewing and consolidating this vast amount of information remains a significant challenge

The advent of large language models (LLMs), which excel at processing and extracting insights from extensive textual data, offers a promising solution. However, general-purpose LLMs like OpenAI’s ChatGPT, Google’s Gemini, or Ought’s Elicit designed for mining literature, present concerns such as their lack of domain specificity, potential for generating inaccurate or incomplete interpretations, and inability to effectively integrate structured scientific databases into their outputs. To address these limitations, we present EpitopeMiner, a domain-specific LLM integrated with a Retrieval-Augmented Generation (RAG) framework(15-17), tailored for identifying immunogenic epitopes from in-silico predicted candidates. By leveraging a custom epitope database, EpitopeMiner combines the retrieval of relevant knowledge with LLM-generated insights, enabling the efficient use of existing data, facilitating cross-study information sharing, and advancing the development of personalized cancer vaccines.

## Methods

### Conventional Workflow for Epitope Prediction

Whole-genome sequencing was performed for a tumor-normal paired from a primary refractory Natural-killer/T cell lymphoma (ENKL) patient. BWA (v0.7.17) was used to aligned the sequencing data to the hs37d5 reference genome and Sambamba (v0.6.5) was used to mark out PCR duplicates. The tumoral and normal data was aligned with an effective coverage of 58.8x and 25.6x, respectively. Strelka (v2.9.10) was performed on the tumor-normal pair aligned BAM files simultaneously and annotation of the candidate variants were annotated by wAnnovar (Jan 2024). Minimum variant-depth of 3x and minimum variant-allele frequency of 5% was used as cut-off to control quality of candidate variants. A final list of 43 non-silent protein-changing variants were curated. The flanking 8 to 11 amino acids of each variant were then tested for its binding affinity to HLA class I molecules.

HLA-typing was performed with Optitype (v1.3.5) on the sequencing data from both tumoral and normal tissues for concordance. The whole-genome data was preprocessed by aligning to the “hla_reference_dna.fasta” downloaded from Optitype’s website, whereby only mappable sequencing reads were curated, for the actual Optitype pipeline, for the final HLA-typing of each sample.

These mutated peptides were analyzed for their ability to bind HLA class I molecules using NetMHCpan v4.1 and MHCflurry v2.0.6. NetMHCpan used Artificial Neural Network (ANNs) to calculate peptide binding affinities of each mutated peptide to a wide range of HLA alleles, providing binding scores (IC50 values) and binding rank percentiles. MHCflurry provided complementary analysis by calculating ligand presentation scores, integrating both peptide binding affinity and surface presentation likelihood. Predictions from NetMHCpan and MHCflurry were combined to select the most promising epitopes. Mutated peptides with a binding affinity threshold of IC50 < 500nM were shortlisted as high-affinity neoantigens. From this analysis, 55 neoantigens were identified. After removing duplicates, 43 unique neoantigen remained. These 43 neoantigens were used to evaluate the performance on epitopes that are not well documented in the existing databases, due to the nature of being highly patient specific.

### Custom Epitope Database

Leveraging the RAG-LLM approach, which combines a retrieval system with a generative AI model, we built a custom database containing MHC-I associated epitope information. This approach allows for efficient extraction and summarization of relevant data from indexed papers. By connecting RAG with an LLM, we can generate contextually rich and accurate summaries, providing insights into epitope characteristics. This integration bridges data retrieval and interpretation, enabling a more comprehensive analysis of MHC-I associated epitopes for immunological studies. Building the database involves three key components: data curation, data indexing, and embedding generation.

#### Data Curation

A total of 3,476 PubMed Identifiers (PMIDs) for all MHC class I T-cell epitopes deposited in the IEDB database were retrieved. Of these, full texts for 1,959 articles were accessible via PubMed Central (PMC) Open Access using the BioC API in JSON format. Figures, captions, tables, and references were removed during preprocessing.

The processed full texts were split into smaller units, referred to as text chunks, to ensure effective retrieval. This was achieved using a hierarchical chunking approach with the *RecursiveCharacterTextSplitter* function from LangChain. Unlike simpler methods, such as the *CharacterTextSplitter*, which splits text based on specific characters and may produce inconsistent chunk sizes, the *RecursiveCharacterTextSplitter* breaks text down from larger units (e.g., paragraphs) into progressively smaller units (e.g., sentences or phrases) while maintaining semantic coherence. This method ensures that the resulting chunks are uniform in size and contextually meaningful, making it well-suited to our use case(18)

Each chunk was configured to contain 1,000 tokens with a 200-token overlap between successive chunks. This setup was chosen to balance providing sufficient context within each chunk and keeping the size manageable for efficient processing.

#### Embedding Generation

To enable efficient similarity search, such as identifying matching epitopes and retrieving relevant information, each text chunk is transformed into a high-dimensional vector representation using advanced natural language processing models. We employ a hybrid search strategy to leverage the complementary strengths of dense and sparse embeddings. This approach combines the semantic depth of dense embeddings with the precision of sparse embeddings, addressing the dual need for contextual understanding and keyword-specific lookups.

Dense embeddings, generated using the text-embedding-3-large model(19), are adept at capturing nuanced semantic relationships, enabling the system to compare and understand text based on meaning. Sparse embeddings, created using SPLADE (Sparse Lexical and Expansion Model for First Stage Ranking)(20), excel at keyword-based searches, which are particularly valuable for finding matching epitopes and related information. By integrating dense embeddings for semantic understanding and sparse embeddings for precise keyword matching, the hybrid system enhances the efficiency of searches, improves relevance scoring, and optimizes storage for large-scale epitope databases.

#### Data Indexing

Once generated, the embeddings are indexed into a specialized vector database, Pinecone(21), which organizes them into a searchable structure, enabling rapid retrieval of relevant embeddings during a query. Among several vector database solutions, such as FAISS (Facebook AI Similarity Search)(22) and ChromaDB(23), Pinecone was selected for its robust support for hybrid search, combining dense and sparse embeddings to enhance both semantic understanding and keyword-specific lookups.

#### Data Retrieval

When a query, either an epitope sequence or free-text, is submitted, it is first converted into an embedding. The system then searches the indexed embeddings to identify the most semantically similar vectors to the query, ensuring accurate and contextually relevant matches.

### EpitopeMiner Workflow

#### User Input

EpitopeMiner processes one epitope sequence at a time. For each sequence, it formulates a query using a predefined template: *“Give information regarding* {*epitope*} *T-cell response*, {*epitope*} *immunogenicity for the candidate epitope* {*epitope*}”; where {*epitope*} is replaced by the user-provided epitope sequence.

#### Similar Epitope Search

To expand the search space in the custom database and retrieve information on epitopes with similar but not identical sequences, the user-provided epitope sequence {*epitope*} is compared against a reference list of MHC-I epitope sequences from IEDB using the Smith-Waterman local alignment method (PairwiseAligner module Biopython, version 1.84). Unlike global alignment methods, this approach focuses on aligning the most similar subsequences within two larger sequences, emphasizing regions of high similarity. Penalties are applied for mismatches (mutations) and gaps (insertions or deletions), reducing the overall alignment score. The 10 most similar epitopes, determined by the highest similarity scores, are then included for processing using the same template query, *“Give information regarding* {*epitope*} *T-cell response*, {*epitope*} *immunogenicity for the candidate epitope* {*epitope*}”.

The rationale behind this approach is that epitopes with similar sequences often share binding affinities to MHC molecules and T-cell receptors, which are essential for eliciting immune responses. This capability is particularly critical for studying neoantigens, which are patient-specific tumor mutations and are less likely to be represented in public datasets. By retrieving information about similar epitopes, EpitopeMiner overcomes the challenge of limited neoantigen representation, enhances the utilization of public data, and creates synergies across studies, ultimately supporting personalized cancer vaccine research.

#### Context Generation

Each of the 11 identified queries (comprising the original epitope and its 10 most similar epitopes), structured as, *“Give information regarding* {*epitope*} *T-cell response*, {*epitope*} *immunogenicity for the candidate epitope* {*epitope*}”, undergoes dense and sparse embedding generation followed by retrieval of the 20 most relevant text chunks from the custom epitope database. This retrieval is performed using a hybrid search approach based on similarity scores (see Methods – Custom Epitope Database).

#### Response Generation

For each of the 11 epitopes (including the original epitope and its 10 most similar epitopes), the corresponding query, *“Give information regarding* {*epitope*} *T-cell response*, {*epitope*} *immunogenicity for the candidate epitope* {*epitope*}” along with the 20 most relevant text chunks retrieved from the custom epitope database, is processed using the GPT-4o API LLM model (OpenAI, San Francisco, CA, USA). To enhance reliability and robustness, five parallel LLM responses are generated for each epitope. These responses are then consolidated into a single consensus output, specifically summarizing information relevant to T-cell response and immunogenicity domains.

#### Structured Responses for Multi-Epitope Queries

To facilitate querying multiple epitopes, EpitopeMiner employs a rule-based JSON structure to format GPT-4o API LLM responses into a tabular format (Figure 2). The resulting table includes epitope sequences, identifiers for associated text chunks (enabling retrieval from the indexed custom database), various LLM parameters, and unique paper counts linked to the defined knowledge domains, such as T-cell response and immunogenicity.

**Figure 1.**
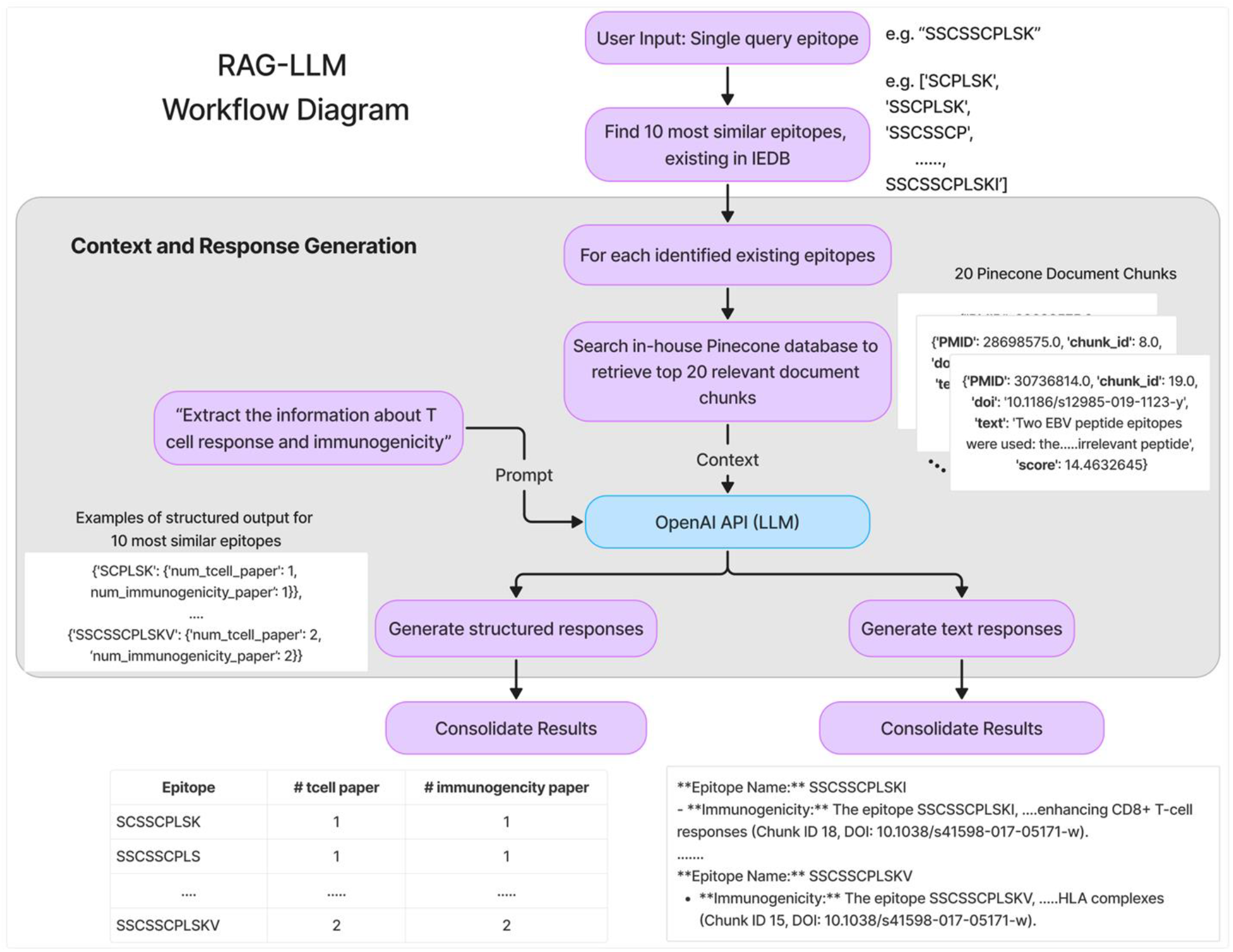
RAG-LLM workflow. The process begins with a user-provided epitope, followed by the identification of 10 similar epitopes. Relevant content for these 11 epitopes is retrieved from an in-house database stored in Pinecone, processed using OpenAI’s LLM API, and presented as structured and textual outputs.

**Figure 2.**
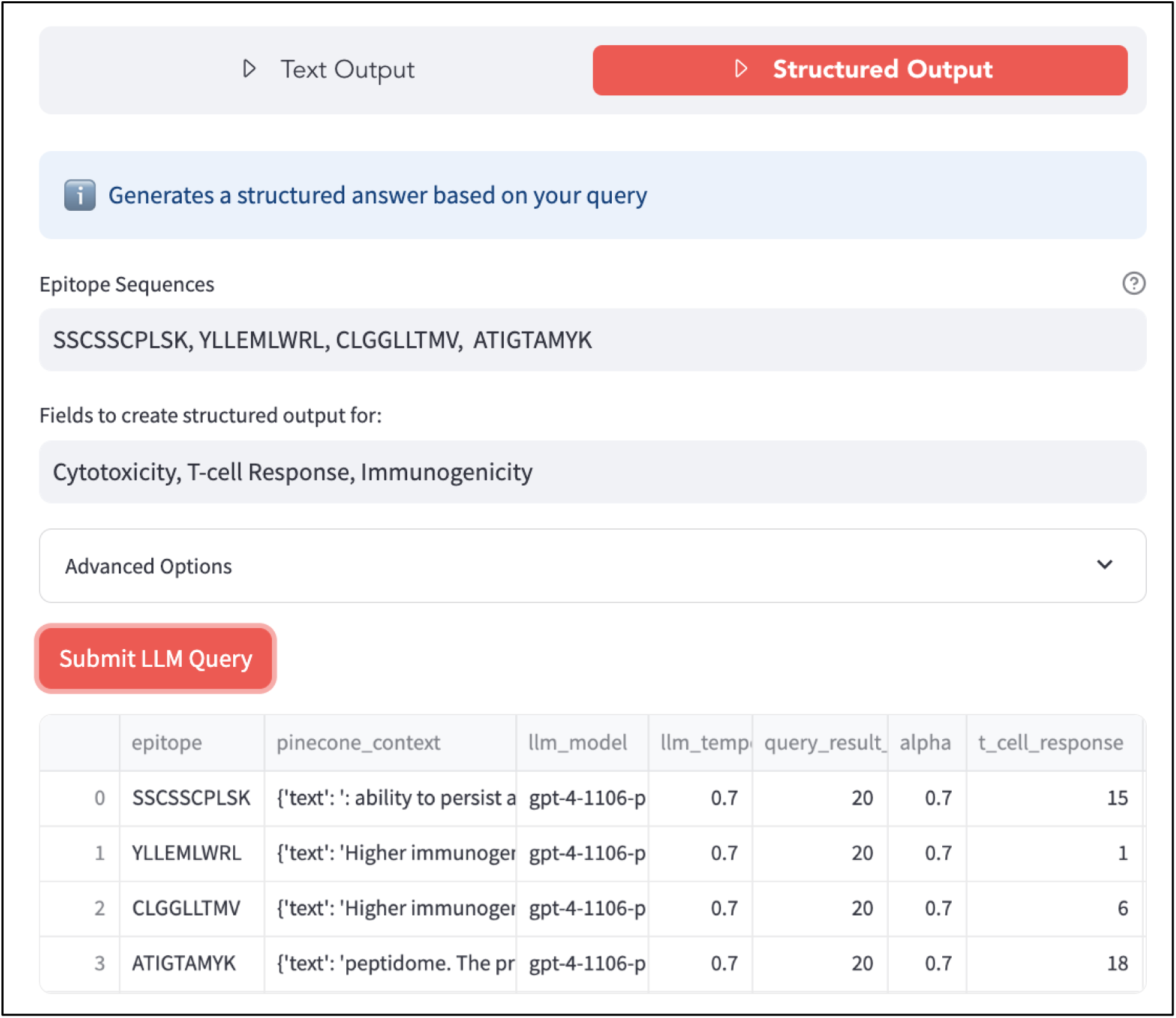
Example of structured responses generated by EpitopeMiner. The output table includes the epitope sequence, associated text chunk identifiers, LLM parameters (e.g., temperature, number of text chunks, alpha value for hybrid search), and the unique number of papers linked to T-cell response.

### Benchmarking Tool - Elicit

Elicit, is an AI-powered research assistant developed by Ought(24), it automates the process of retrieving, summarising and also provides a free-text querying function allowing information tabulation using custom column creation function. Elicit has indexed over 125 million research papers across a wide range of disciplines built on the Semantic Scholar corpus.

Given a user-input free-text description, Elicit generates a summary highlighting information relevant to the user query, using the 8 most relevant (or recent) papers. It also allows the creation of custom columns to identify specific information about each paper using AI. For benchmarking against EpitopeMiner, we defined two new columns: one based on the query, “Does the paper mention T-cell response?” and the other, “Does the paper mention immunogenicity?”. Additionally, we expanded the search to include the top 16 most relevant papers.

### ChatGPT Usage for Manuscript Preparation

ChatGPT (OpenAI, San Francisco, CA, USA) was used to assist in drafting and refining sections of this manuscript, including language optimization and style consistency. The content was thoroughly reviewed and validated by the authors to ensure accuracy and intellectual integrity.

## Results

EpitopeMiner processes queried epitopes individually. Unlike general-purpose LLM tools, it first searches a custom epitope database for T-cell response-related information, ensuring domain-specific queries are refined before being processed by the OpenAI API. Furthermore, EpitopeMiner provides summaries tailored to specific knowledge domains, currently focused on T-cell response and immunogenicity, and includes the associated text chunks retrieved from the epitope database to facilitate interpretation and validation. We evaluate the performance of EpitopeMiner using well-characterized MHC class I epitopes from the IEDB database, as well as lymphoma-derived neoantigens, which are highly patient-specific and minimally documented.

### EpitopeMiner Evaluation with Well-Characterized IEDB Epitopes

Thirteen well-characterized MHC class I epitopes from the IEDB database were randomly selected to evaluate EpitopeMiner’s performance, benchmarked against Elicit, a commercial tool designed for literature review. Table 1 summarizes the structured output generated by EpitopeMiner for the 13 epitopes, alongside paper counts returned by Elicit using its custom column creation feature. Standard queries, such as “Does the paper mention T-cell response?” or “Does the paper mention immunogenicity?”, were applied to the 16 most relevant research articles retrieved by Elicit.

**Table 1.**
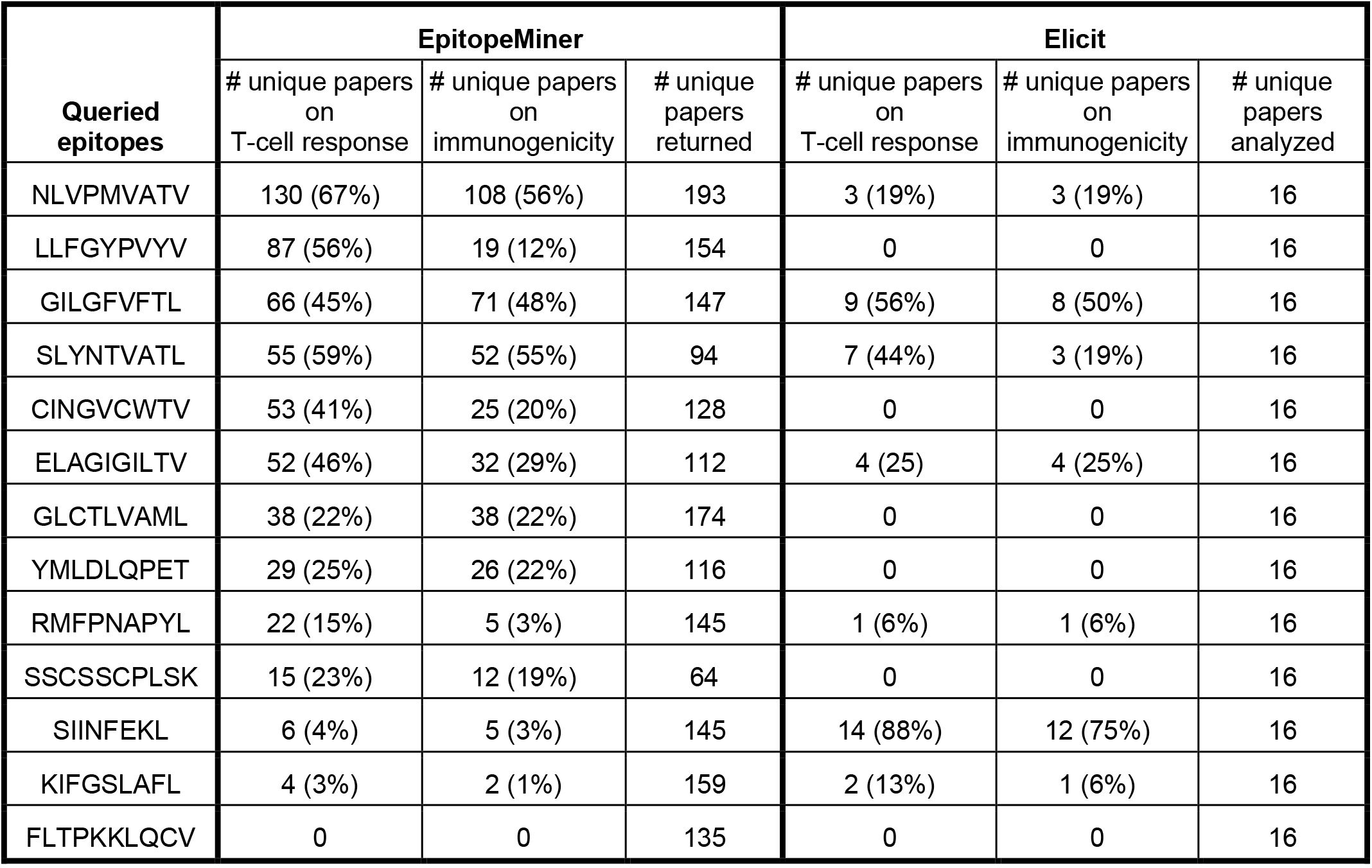
RAG-LLM Performance Evaluation Using 13 Randomly Selected Epitopes from IEDB Database, Benchmarked Against Elicit. The RAG-LLM results include the queried epitope along with its 10 most similar epitopes. For comparison, paper counts from Elicit’s custom column creation feature, querying “T-cell response” or “immunogenicity” in the top 16 most relevant research papers, are also listed. Percentage values indicate the proportion of papers among all unique papers returned or analyzed. Rows are ordered by the decreasing number of papers associated with T-cell response and immunogenicity identified by RAG-LLM.

Assuming that epitopes with similar sequences share binding affinities to MHC molecules and T-cell receptors for eliciting immune responses, EpitopeMiner identifies the 10 most similar epitopes from the custom database and generates responses for each, along with the original queried epitope. Among the 13 queried epitopes, EpitopeMiner retrieved information, in terms of papers in the custom database, for 12 of them, while Elicit found relevant papers for 7 epitopes. Furthermore, EpitopeMiner consistently retrieves a higher percentage of papers associated with T-cell response and immunogenicity, with the exception of SIINFEKL. For well-established epitopes like SIINFEKL, access to a larger database, such as the one utilized by Elicit, proves advantageous. Notably, a manual evaluation of the 16 unique papers retrieved by Elicit reveals relevant and valuable information about the T-cell response and immunogenicity of the target epitope.

### EpitopeMiner Evaluation with Lymphoma-Derived Neoantigens

Neoantigens are novel peptides arising from tumor-specific mutations and are more likely to elicit a robust T-cell-mediated immune response, making them critical targets for personalized cancer vaccine development. However, limited existing knowledge poses a challenge. By leveraging EpitopeMiner’s similar epitope search capability, we retrieved relevant literature information for 43 neoantigen epitopes (MHC-I) identified from a lymphoma patient through conventional prediction tools (see Methods).

Table 2 shows that EpitopeMiner identified similar epitopes for 8 out of the 43 neoantigen epitopes, with each epitope having references in nearly 150 papers indexed in the custom database. Conversely, paper counts obtained from Elicit’s custom column creation feature returned no relevant papers. Among the papers retrieved by EpitopeMiner, a smaller proportion is specific to T-cell response and immunogenicity compared to the well-characterized IEDB epitopes, as shown in Table 1.

**Table 2.**
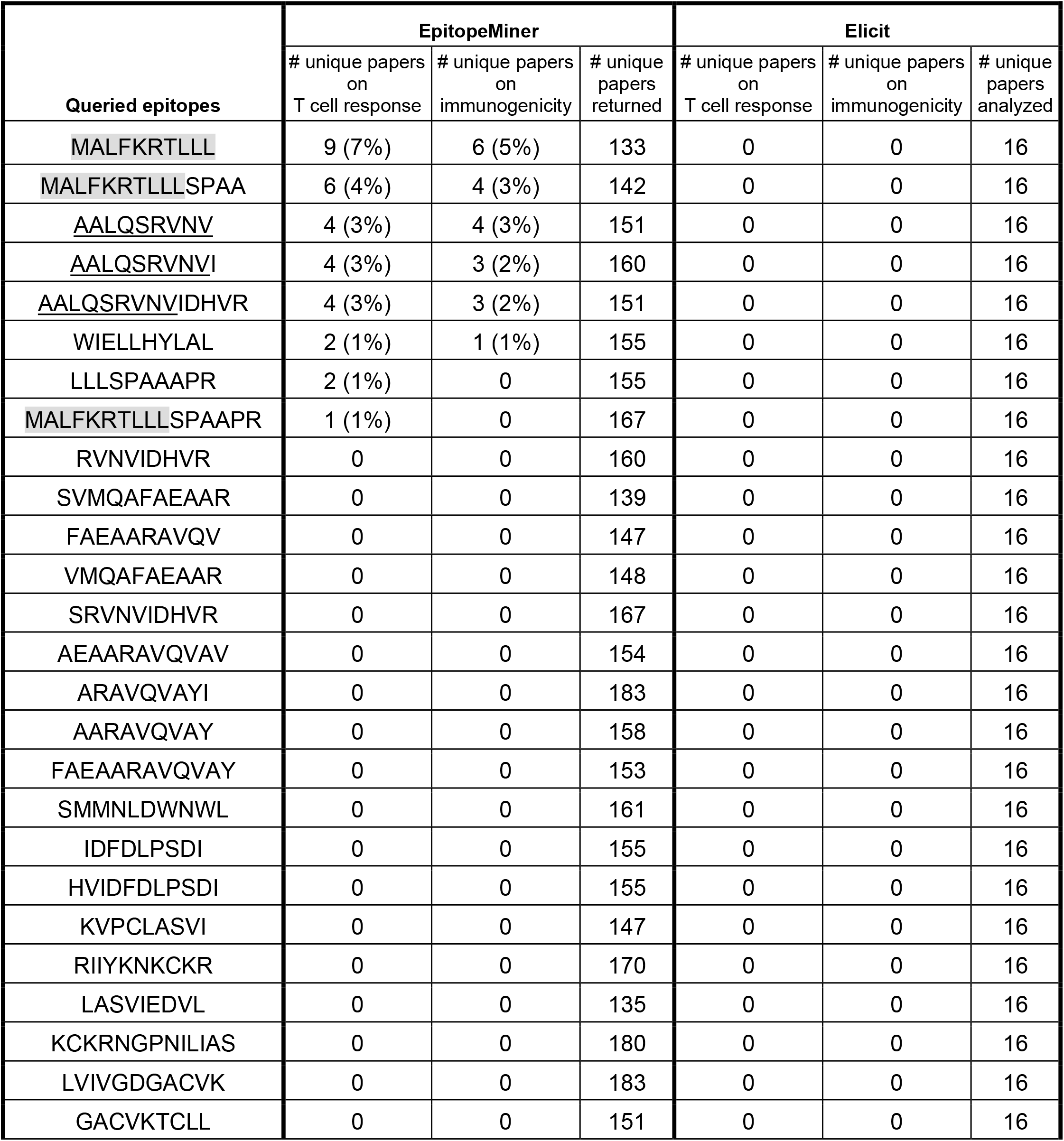
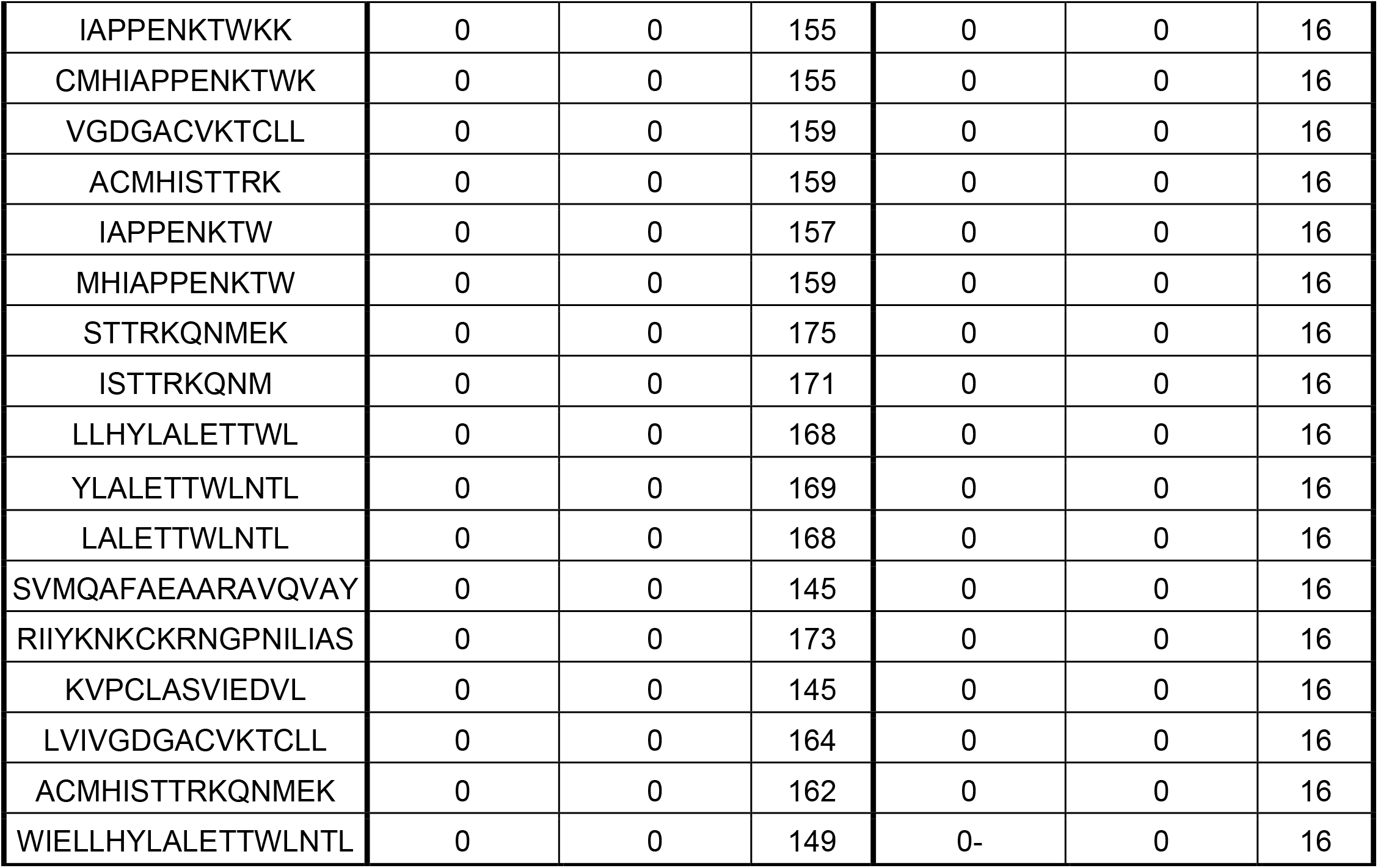
RAG-LLM Performance Evaluation Using 43 Neoantigens from a Lymphoma Patient, Benchmarked Against Elicit. The RAG-LLM results include the queried epitope along with its 10 most similar epitopes. For comparison, paper counts from Elicit’s custom column creation feature, querying “T-cell response” or “immunogenicity” in the top 16 most relevant research papers, are also listed. Percentage values indicate the proportion of papers among all unique papers returned or analyzed. Rows are ordered by the decreasing number of papers associated with T-cell response and immunogenicity identified by RAG-LLM. Three epitopes containing the sequence ‘MALFKRTLLL’ and another three containing the sequence ‘AALQSRVNV’ were underlined and shaded (see Tables 3 and 4 for detailed analysis).

Among the top-ranked neoantigen epitopes based on the unique number of papers associated with T-cell response and immunogenicity, some share common sequences. Specifically, ‘MALFKRTLLL’ is a sequence present in ‘MALFKRTLLL’ ‘MALFKRTLLLSPAA’ and ‘MALFKRTLLLSPAAPR,’ while ‘AALQSRVNV’ is shared among ‘AALQSRVNV’ ‘AALQSRVNVIDHVR’ and ‘MALFKRTLLLSPAAPR (Table 2). Table 3 presents EpitopeMiner’s detailed responses for the queried neoantigen epitope, MALFKRTLLL. Among the 10 most similar epitopes identified, 3 were referenced in papers indexed within the custom database, and they either comprise “VMAPRTLLL” or a similar sequence, such as “VMPPRTLLL” It is worth noting that no direct reference was found for the queried epitope. The textual responses provided include information such as HLA restrictions, peptide family, and associations with viruses like Human Cytomegalovirus (HCMV).

**Table 3.**
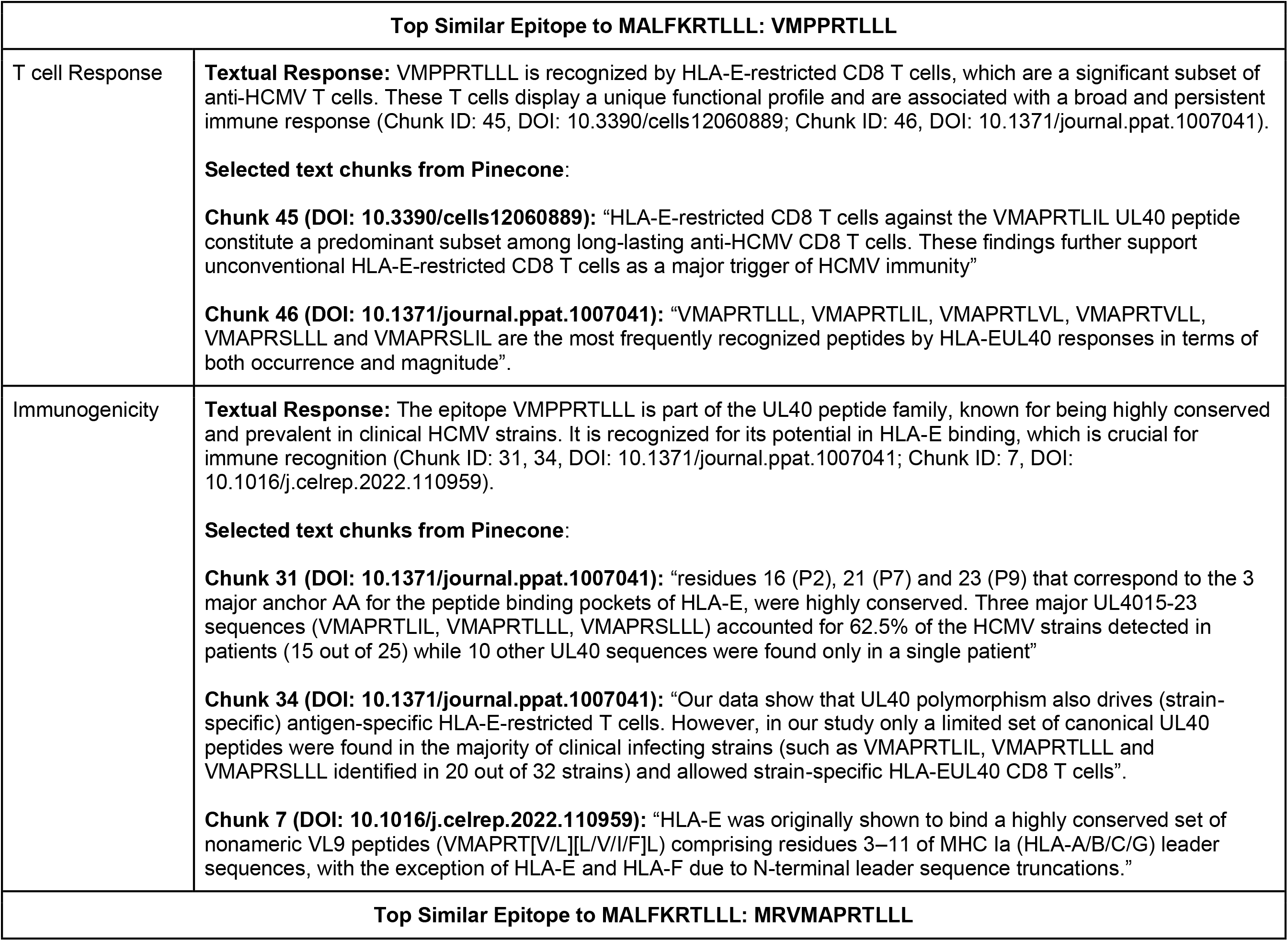
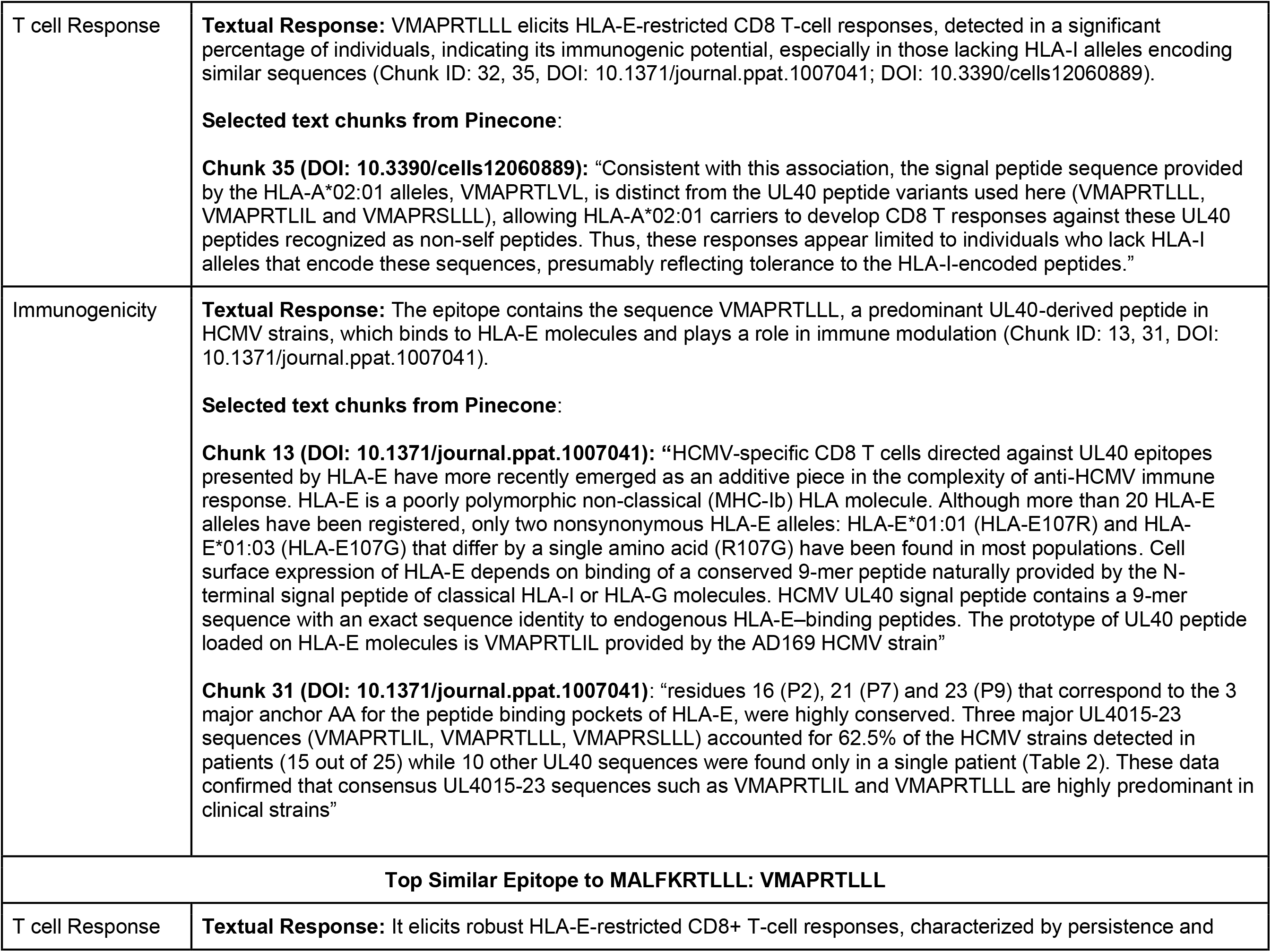
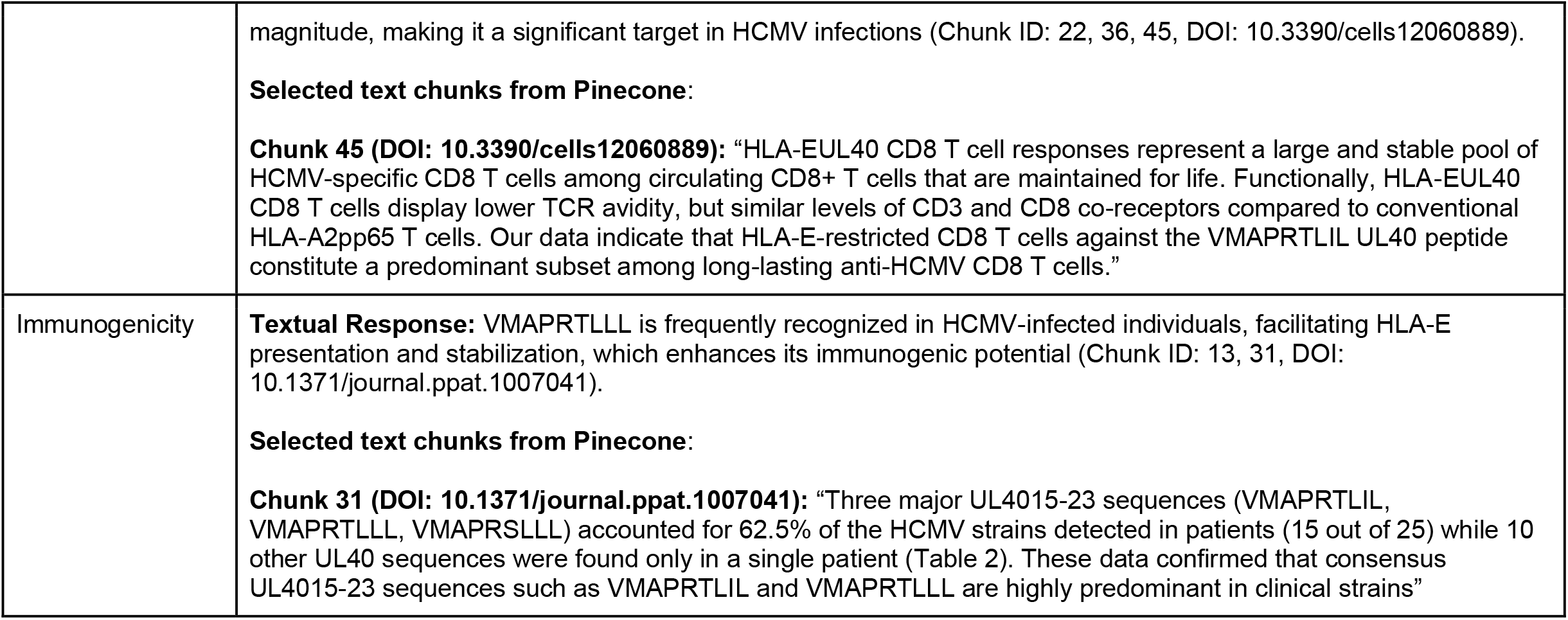
Detailed Responses Generated by the RAG-LLM Pipeline for the Queried Epitope MALFKRTLLL and Its 10 Most Similar Epitopes. Among the 11 epitopes analyzed, no information was retrieved for the queried epitope itself or 7 of its most similar epitopes. For the remaining 3 similar epitopes, the pipeline provided textual responses and selected text excerpts from associated journal articles indexed in the Pinecone database.

To support validation and knowledge expansion, EpitopeMiner also returns text chunks and paper DOIs from the indexed custom database (Pinecone) to users, providing context for downstream processing by the OpenAI API. Due to space constraints, selected text chunks are presented in Table 3. Inspection of the original text chunks reveals that the OpenAI LLM API incorporated additional similar epitope searches, effectively expanding the search space. Three of the ten most similar epitopes were referenced in papers indexed in the custom database, despite lacking direct references. It is also noteworthy that no direct references were found for the queried epitope. Notably, EpitopeMiner’s textual responses provided additional insights into epitope characteristics, including structural details and T-cell receptor (TCR) information. For the selected neoantigen epitope ‘AALQSRVNV’ (Table 4), similar findings were observed. Three of the ten most similar epitopes were referenced in papers indexed in the custom database, despite lacking direct references. It is also noteworthy that no direct references were found for the queried epitope. Notably, EpitopeMiner’s textual responses provided additional insights into epitope characteristics, including structural details and T-cell receptor (TCR) information.

**Table 4.** Detailed Responses Generated by the RAG-LLM Pipeline for the Queried Epitope AALQSRVNV and Its 10 Most Similar Epitopes. Among the 11 epitopes analyzed, no information was retrieved for the queried epitope itself or 7 of its most similar epitopes. For the remaining 3 similar epitopes, the pipeline provided textual responses and selected text excerpts from associated journal articles indexed in the Pinecone database.

| Top Similar Epitope to AALQSRVNV: QAALQSAWQ |  |
| --- | --- |
| T cell Response | **Textual Response:** Similar epitopes, such as QW9, elicit robust CD8+ T-cell responses when presented by HLA-B*5801, suggesting QAALQSAWQ might also elicit comparable responses if structurally similar (Chunk ID: 4, DOI: 10.1007/s13238-016-0309-y).<br>**Pinecone Context (selected):**<br>**Chunk 4 (DOI: 10.1007/s13238-016-0309-y):** “To begin to address the structural basis for T cell mediated control of HIV, we initiated a structural analysis of an immunodominant CD8+ T cell epitope QW9 (QASQEVKNW, p24gag residues 176–184) from HIV-1 bound to the restricting HLA class I molecule B*5801. This epitope is of particular interest because the conserved Gag protein forms the essential capsid of HIV-1 (Zhao et al.), with mutations likely to impose a fitness cost”. |
| Immunogenicity | **Textual Response:** The QAALQSAWQ epitope does not have direct information available. However, similar epitopes like QW9 (QASQEVKNW) show strong immunogenic potential due to stable interaction with HLA-B*5801, indicating potential immunogenicity for QAALQSAWQ if it shares similar structural features (Chunk ID: 4, DOI: 10.1007/s13238-016-0309-y).<br>**Pinecone Context (selected):**<br>**Chunk 4 (DOI: 10.1007/s13238-016-0309-y):** “To begin to address the structural basis for T cell mediated control of HIV, we initiated a structural analysis of an immunodominant CD8+ T cell epitope QW9 (QASQEVKNW, p24gag residues 176–184) from HIV-1 bound to the restricting HLA class I molecule B*5801. This epitope is of particular interest because the conserved Gag protein forms the essential capsid of HIV-1 (Zhao et al.), with mutations likely to impose a fitness cost”. |
| **Top Similar Epitope to AALQSRVNV: YFLEAALQSAY** |  |
| T cell Response | **Textual Response:** Not sufficient information to analyze T-cell response for this epitope directly. YLQ elicits strong CD8+ T-cell responses with public TCR recognition, indicating potential for similar responses in YFLEAALQSAY if it shares structural features (Chunk ID: 50.0, DOI: 10.1038/s41467-021-27669-8).<br>**Pinecone Context (selected):**<br>**Chunk 50 (DOI: 10.1038/s41467-021-27669-8):** “We found that two natural variants of the YLQ and RLQ epitopes (P272L and T1006I) that contain the most commons epitope mutations in the GISAID database had different effect on TCR recognition. Despite 1.3- to 6.8-fold reductions in TCR affinity as measured by SPR, T1006I activated RLQ3, RLQ5, RLQ7, and RLQ8 T cells in functional assays nearly as efficiently as the wild-type epitope. Since the RLQ response is oligoclonal, the efficiency of recognition of the T1006I variant could vary among patients depending on the involvement of particular clones” |
| Immunogenicity | **Textual Response:** Not sufficient information to analyze immunogenicity for this epitope directly. The similar epitope YLQPRTFLL (YLQ) is highly immunogenic with strong binding affinity to HLA-A*02:01 (Chunk ID: 9.0, DOI: 10.1038/s41467-021-27669-8).<br>**Pinecone Context (selected):**<br>**Chunk 9 (DOI: 10.1038/s41467-021-27669-8):** “The YLQ epitope has been identified as immunodominant in multiple independent studies, including one involving the BNT162b2 mRNA vaccine. For its part, the RLQ epitope, unlike YLQ, is conserved across human and zoonotic sarbecoviruses and is therefore a potential candidate for |
|  | inclusion in a pan-sarbecovirus vaccine.” |
| **Top Similar Epitope to AALQSRVNV: VPHLEAALQEY** |  |
| T cell Response | **Textual Response:** Similar epitopes elicit robust CD8+ T-cell responses, indicating VPHLEAALQEY may also elicit comparable responses (Chunk ID: 39, DOI: 10.1186/1475-2875-12-185).<br>**Pinecone Context (selected):**<br>**Chunk 39 (DOI: 10.1186/1475-2875-12-185):** "E2(FVEALFQEY): This sequence was supported as a B35:01-restricted epitope (B07 supertype) as the synthesized peptide was active with v69 (Table 5). E2 was also predicted by NetMHC to be an A01:01-restricted epitope (A01 supertype) using v05; however, PBMC were not available from v05 to test the synthesized peptide." |
| Immunogenicity | **Textual Response:** Similar epitopes like FVEALFQEY demonstrate broad immunogenic potential across different HLA supertypes, suggesting similar potential for VPHLEAALQEY (Chunk ID: 39, DOI: 10.1186/1475-2875-12-185).<br>**Pinecone Context (selected):**<br>**Chunk 39 (DOI: 10.1186/1475-2875-12-185):** "E2(FVEALFQEY): This sequence was supported as a B35:01-restricted epitope (B07 supertype) as the synthesized peptide was active with v69 (Table 5). E2 was also predicted by NetMHC to be an A01:01-restricted epitope (A01 supertype) using v05; however, PBMC were not available from v05 to test the synthesized peptide. E2, however, could be tested with v127, also A01:01-restricted, and was active. Therefore E2 appears restricted by two allele groups, B35:01 and A01:01 that belong to different supertypes, B07 and A01, respectively."* |

## Discussion

Personalized cancer vaccines hold significant promise for inducing specific and robust immune responses, leading to effective tumor clearance. However, the current vaccine development pipeline remains time-consuming and costly, primarily because conventional prediction methods focus heavily on MHC molecule binding affinity. In contrast, effective immune responses depend on multiple factors, particularly T-cell response and immunogenicity, with an expanding list of contributing elements. Consequently, significant time and effort are required to validate the extensive list of epitope candidates generated by prediction tools, either through experimental methods or labor-intensive literature reviews.

To address this, we present EpitopeMiner, a domain-specific language model designed to identify immunogenic epitopes by utilizing a custom database of epitope-related full-text papers. The application of EpitopeMiner to well-characterized epitopes from IEDB demonstrates its ability to consistently return relevant papers, efficiently retrieving targeted information, including insights on T-cell response and immunogenicity. In contrast, Elicit provides information for only a subset of the tested epitopes, despite accessing a larger database. This highlights the advantage of EpitopeMiner’s context-enriched LLM search in enhancing both specificity and coverage.

However, retrieving specific and comprehensive information about neoantigens remains challenging, as they are patient-specific tumor mutations and are less likely to be represented in public datasets. In this context, the ability to identify epitopes with similar sequences to the queried neoantigen epitopes becomes crucial. Hence, EpitopeMiner integrates a similar epitope search function, designed to identify reported epitopes with comparable sequences and potentially similar immunogenicity. While Elicit fails to retrieve relevant papers, EpitopeMiner identifies papers for similar sequence epitopes, facilitating knowledge extraction from public data and creating synergies across studies, thereby accelerating cancer vaccine development research.

Two other key features of EpitopeMiner to highlight are its structured output for each engineered knowledge domain, currently focusing on T-cell response and immunogenicity, and its ability to extract and interpret original text chunks and papers directly from its database. In contrast, while Elicit provides access to papers, it requires manual searching for specific epitopes. Although Elicit offers a search and summarization feature through custom column creation, it yields significantly fewer unique papers related to T-cell response and immunogenicity when analyzing well-characterized IEDB epitopes and retrieves zero papers in the lymphoma-derived neoantigen analysis.

Our study has several limitations. First, our custom database is relatively small, comprising approximately 1.9 thousand full-text papers. We aim to expand this database by incorporating more publications and integrating structured public databases, particularly those focusing on T-cell response information, such as dbPepNeo(13) and NeoDB(14). Additionally, the database is currently limited to MHC-I associated epitopes, while the significance of MHC-II associated immune responses is increasingly recognized and should be accounted for in future studies. Second, EpitopeMiner is currently limited to two specific knowledge domains: T-cell response and immunogenicity. However, the modular design of the RAG-LLM framework allows for the addition of new domains in the future. Other factors influencing effective immune responses by epitopes, such as peptide stability, processing and presentation efficiency, and the role of HLA allotypes, should be considered in future iterations. Third, further validation is needed using patient tumor-derived neoantigens from diverse tumor types to ensure broader applicability and robustness. Fourth, unlike Elicit, EpitopeMiner is not intended for general-purpose literature searches and requires a specific epitope sequence for queries. Additionally, it does not currently support iterative or narrowed-down information retrieval.

In summary, EpitopeMiner offers a novel and effective approach for retrieving comprehensive and targeted information related to queried epitopes. Specifically designed to extract knowledge relevant to tumor-specific neoantigens, it leverages similar sequence epitopes to enhance data accessibility. By integrating EpitopeMiner with conventional prediction tools, we anticipate significant improvements in the efficiency of personalized vaccine development, reducing time requirements and enabling cost-effective shortlisting of high-potential epitope candidates.

